# Control of coronary lymphangiogenesis by epicardial VEGFC/D

**DOI:** 10.1101/2023.02.10.528007

**Authors:** Ester de la Cruz, Vanessa Cadenas, Susana Temiño, Guillermo Oliver, Miguel Torres

**Affiliations:** Cardiovascular Regeneration Program, Centro Nacional de Investigaciones Cardiovasculares (CNIC), Madrid, 28029, Spain; Center for Vascular and Developmental Biology, Feinberg Cardiovascular and Renal Research Institute, Northwestern University, Chicago, IL 60611, USA

**Author notes:** Corresponding author **Address:** Centro Nacional de Investigaciones Cardiovasculares. 3, Melchor Fernández Almagro. Madrid 28029. Spain, **Email:**.

## Abstract

The contractile ability of the mammalian heart critically relies on blood coronary circulation, essential to provide oxygen and nutrients to myocardial cells. In addition, the lymphatic vasculature is essential for the myocardial immune response, extracellular fluid homeostasis and response to injury. Recent studies identified different origins of coronary lymphatic endothelial cells, however, the cues that govern coronary lymphangiogenesis remain unknown. Here we show that the coronary lymphatic vasculature develops in intimate contact with the epicardium and with epicardial-derived cells. The epicardium expresses the lymphangiogenic cytokine VEGFC and its conditional elimination from the epicardium abrogates coronary lymphatic vasculature development. Interestingly, VEGFD is also expressed in the epicardium and cooperates with VEGFC in coronary lymphangiogenesis, but it does so only in females, uncovering an unsuspected sex-specific role for this cytokine. These results identify a role for the epicardium/subepicardium as a signalling niche required for coronary lymphangiogenesis and VEGFC/D as essential mediators of this role.

## Introduction

Besides the blood coronary vasculature, the ventricular myocardium also contains a fully developed network of cardiac lymphatics (Brakenhielm & Alitalo, 2019). Coronary lymphatics promote cardiomyocyte proliferation (Liu *et al*, 2020) and are important in the response to myocardial infarction (Henri *et al*, 2016; Klotz *et al*, 2015; Liu *et al*., 2020; Maruyama *et al*, 2021; Vieira *et al*, 2018) congestive heart failure (Witte *et al*, 1969), atherosclerosis (Lim *et al*, 2013; Milasan *et al*, 2016) and heart regeneration in zebrafish (Gancz *et al*, 2019; Harrison *et al*, 2019; Vivien *et al*, 2019). Cardiac lymphatics are first seen along the great arteries and sinus venosus at E12.5-E13.5 (Flaht *et al*, 2012; Karunamuni *et al*, 2010). Most embryonic and cardiac lymphatic endothelial cells (LECs) have a venous origin (Klotz *et al*., 2015; Wigle & Oliver, 1999); however, mesenchymal precursors from the second heart field (SHF) also contribute to ventral coronary LECs (Lioux *et al*, 2020; Maruyama *et al*, 2019). The transcription factor *Prox1* (Wigle & Oliver, 1999), as well as the growth factors VEGFC and VEGFD and their receptor VEGFR3 (Baldwin *et al*, 2005; Bower *et al*, 2017; Haiko *et al*, 2008; Karkkainen *et al*, 2004; Karkkainen *et al*, 2001; Paquet-Fifield *et al*, 2013) are essential during developmental lymphangiogenesis. In the heart, local signalling pathways like Retinoic acid (RA) and macrophage-derived hyaluronan promote cardiac lymphatic vessel maturation and remodelling (Cahill *et al*, 2021; Lioux *et al*., 2020), while the Sema3E-PlexinD1 axis is involved in coronary artery and lymphatic vessel patterning (Maruyama *et al*., 2021). Despite this progress, the cellular and molecular drivers of coronary lymphangiogenesis in the developing heart remain unknown. In several tissues/organs, arteries guide lymphatic vessel growth through VEGFC and CXCL12 expression from endothelial and smooth muscle cells (Cha *et al*, 2012; Vaahtomeri *et al*, 2017). Here, we characterize the growth pattern of coronary lymphatics and report that they do not follow arteries or veins as they colonize the ventricles but rely on intimate interaction with the epicardium and epicardial-derived cells. Expression of VEGFC/D from the epicardium is essential for the lymphangiogenic function of the epicardium.

## Results

To identify the mechanisms that guide coronary lymphangiogenesis, we studied the distribution of cardiac lymphatics as they grow into the ventricles of the mouse fetal heart (Figure 1A-D). Cardiac lymphatics appear at the base of the ventricles, dorsally at the sinus venosus and ventrally on the great arteries around E12.5, however, they do not start colonizing the ventricles until E14.5 (reviewed in (Klaourakis *et al*, 2021)). At E14.5, coronary veins and arteries are already present; however, lymphatic vessels do not follow the pattern of coronary arteries or veins as they colonize the ventricles (Figure 1A-B”, D). In histological sections at E15.5 we observed that coronary lymphatic vessels grow in direct association with the epicardium (Figure 1C-C”). Coronary veins also grow within the subepicardial space, however, when lymphatic vessels coincide with veins in the subepicardium, the lymphatic vessel is always located closer to the epicardium, representing the most superficial coronary vasculature (Figure 1C’, C’’, E). We thus found that coronary lymphatic vessels do not follow arteries or veins, as it happens in other tissues/organs (Cha *et al*., 2012; Vaahtomeri *et al*., 2017), but rather grow freely beneath the epicardium, without following either veins, which grow beneath the lymphatic vasculature, or arteries, which grow within the myocardium. This behaviour correlates with the strong expression of VEGFC and CXCL12 in the epicardium (Cavallero *et al*, 2015; Chen *et al*, 2014).

**Figure 1.**
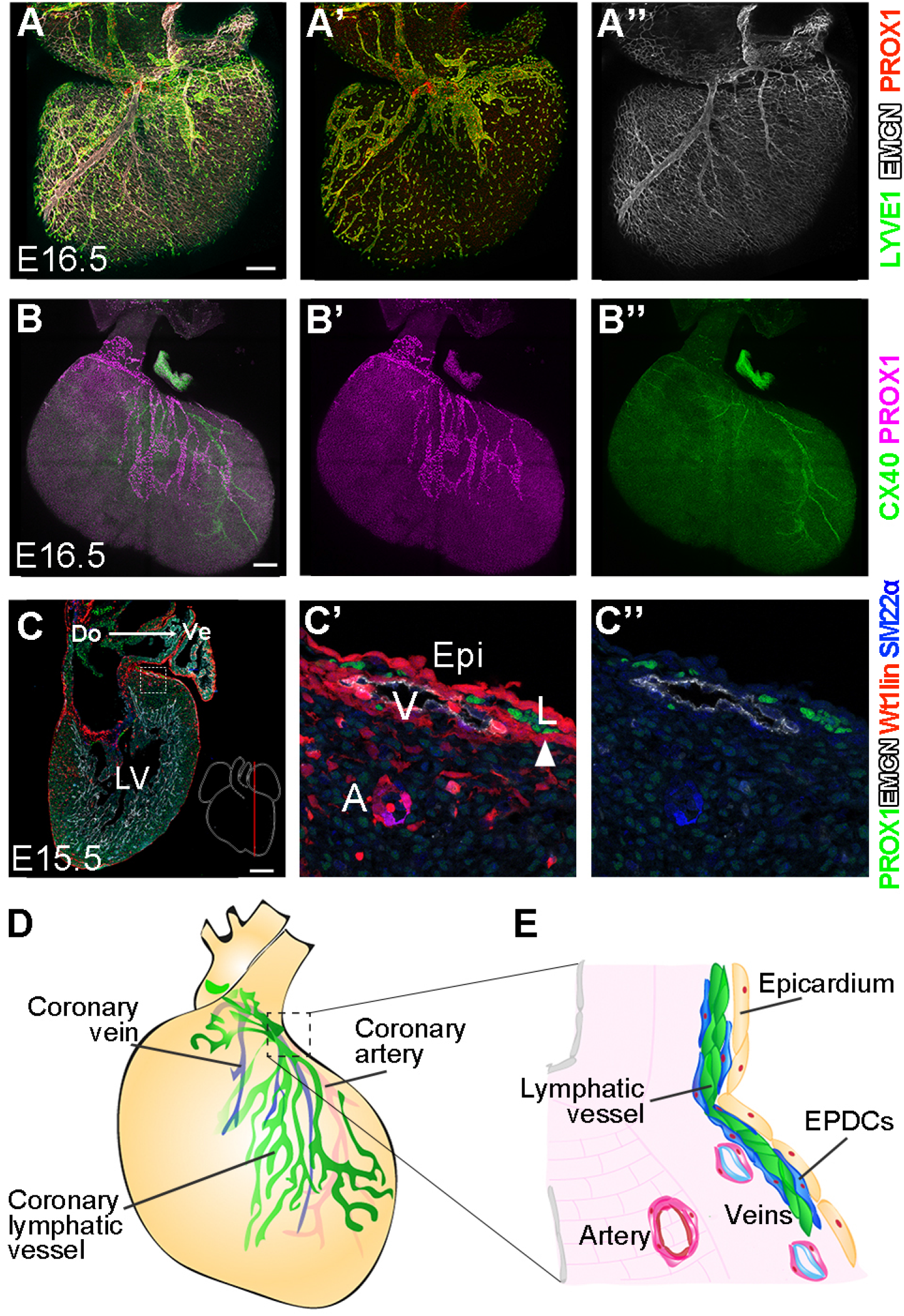
Subepicardial coronary lymphangiogenesis. **(A-A’’**) Detection of veins with Endomucin (EMCN) and lymphatic vessels with Lyve1 and Prox1 in the dorsal side of E16.5 mouse hearts. (**B-B”**). Detection of arteries with Connexin 40 (CX40) and lymphatic vessels with PROX1 in the ventral side of an E16.5 mouse heart. (**C-C’’**) Detection of arteries (A), veins (V) and lymphatic vessels (L) in a sagittal section of an E15.5 *Wt1^Cre^; Rosa26R^Tomato^* mouse heart, which reveals the lineage of cells that express *Wt1^Cre^* (Wt1lin), including the epicardium (Epi). Arrowhead in C’ indicates lymphatic vessel-associated EPDCs. Do: dorsal; Ve: ventral; LV, left ventricle. (**D**, **E**) Schemes of the disposition of coronary vessels in a developing mouse heart. EPDC: epicardial-derived cells.

While VEGFC is strongly expressed in the epicardium (Chen *et al*., 2014), it is also expressed in other cardiac cells (Cahill *et al*., 2021); we therefore specifically studied the relevance of epicardial VEGFC in coronary lymphangiogenesis. We first studied the sensitivity of the developing lymphatic vasculature to increased epicardial VEGFC. For this, we used the *Eef1a1^VegfcGOF^* allele (Pichol-Thievend *et al*, 2018), which provides conditional Cre-mediated VEGFC overexpression, combined with *Wt1^Cre^* (Wessels *et al*, 2012), which drives Cre for activation in the epicardium. Hearts with epicardial VEGFC overexpression showed increased coronary lymphatic coverage (Figure 2A-D), indicating that coronary lymphangiogenesis is sensitive to epicardial VEGFC levels. Next, we studied the requirement for VEGFC function in epicardium/EPDCs, by crossing *Vegfc^flox/flox^* (Lim *et al*, 2019) with *Wt1^Cre^* and analysing the lymphatic vasculature at E16.5. Homozygous mutant hearts (*Vegfc-KO*) showed 50% reduction of lymphatic coverage in the ventral side of the ventricles and 68% reduction in the dorsal side (Figure 2E-H). Total lymphatic length and lymphatic vessel bifurcations were also reduced in *Vegfc-KO* hearts (Figure 2I-L). To confirm these results with an independent Cre line, we induced epicardial-specific deletion of *Vegfc* using *Tbx18^Cre^* (Cai *et al*, 2008). Mutant hearts in this case lacked all lymphatic vessels of the dorsal side of the heart and showed only vestigial lymphatic vessels on the ventral side (Figure 2M-P). These results suggest that the incomplete penetrance observed in the *Wt1^Cre^-* deleted hearts is due to lower efficacy of this Cre line compared with *Tbx18^Cre^*. Dosed levels of epicardial VEGFC are thus essential for coronary lymphangiogenesis, with more relevance in the dorsal than in the ventral side of the ventricles.

**Figure 2.**
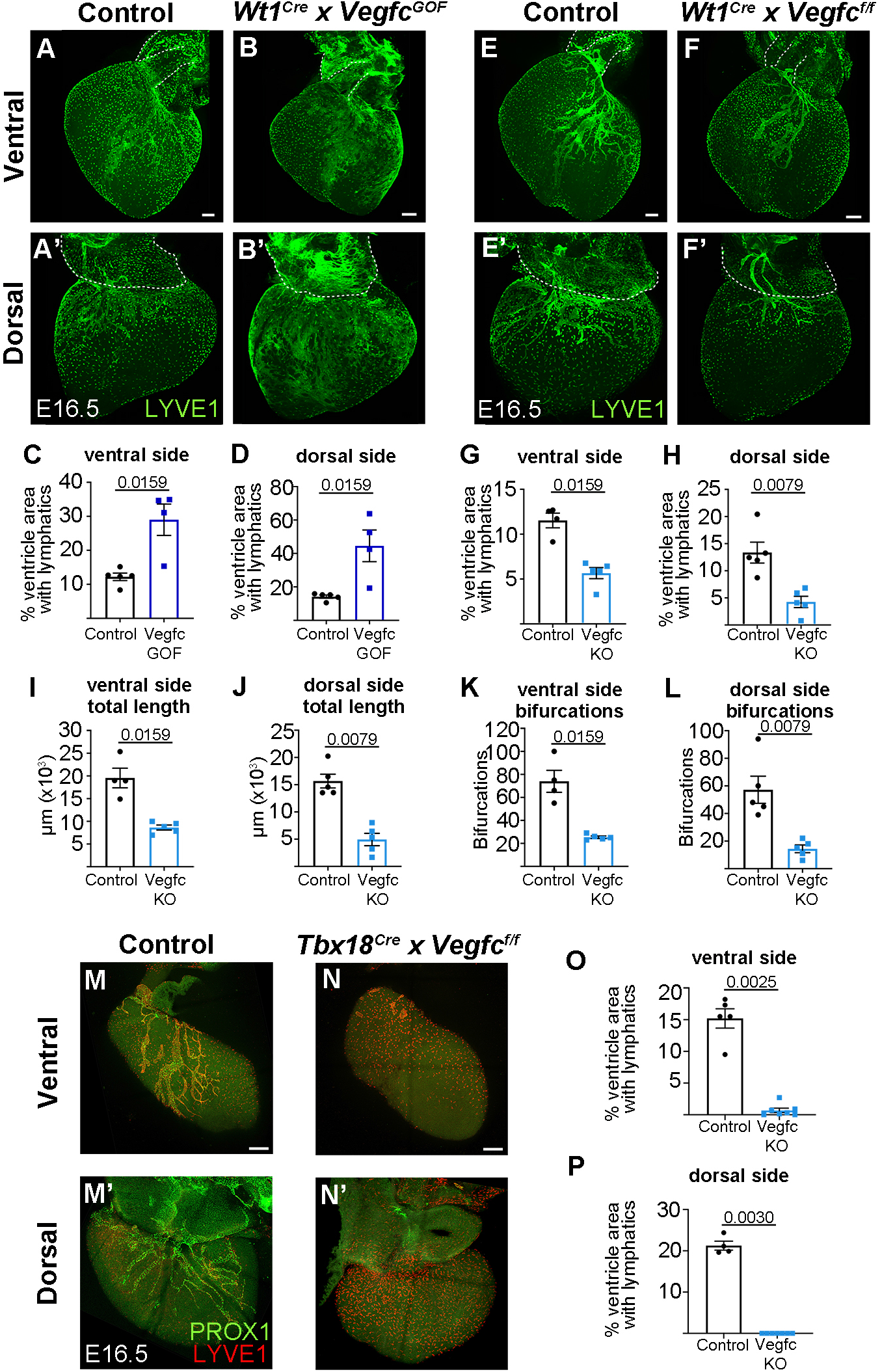
Epicardial VEGFC is essential for coronary lymphangiogenesis. (**A-D**) Representative specimens and quantifications of lymphatic vessel coverage of the dorsal and ventral ventricular surfaces upon *Vegfc* overexpression by activation of *Eef1a1^VegfcGOF^ with Wt1^Cre^ (Vegfc ^GOF^*). (**E-L**) Representative specimens and quantifications of lymphatic vessel coverage, lymphatic length and bifurcations of the dorsal and ventral ventricular surfaces upon *Vegfc* conditional elimination with *Wt1^Cre^*. (**M-P**) Representative specimens and quantifications of lymphatic vessel coverage of the dorsal and ventral ventricular surfaces upon *Vegfc* conditional elimination with *Tbx18^Cre^*. Statistics: Mann-Whitney test with two-tailed p-values.

Although less prominent than VEGFC, VEGFD collaborates with VEGFC in promoting lymphangiogenesis in the developing intestinal lymphatic vasculature (Nurmi *et al*, 2015). Whole-mount detection of VEGFD protein in WT E16.5 hearts revealed expression in the epicardium (Figure 3A, B) and epicardial-derived cells associated to the growing coronary lymphatic vasculature (Figure 3C, C’). To functionally assess the role of VEGFD in coronary lymphatic development, we generated *Vegfd* global knockouts using CRISPR-Cas9 deletion of exons 3 and 4, which codify a portion of VEGFD central receptor-binding domain, *“VEGF homologous domain”* (VHD), essential for its function (Achen *et al*, 1998) (See Methods). As previously described, VEGFD deletion produced viable and fertile animals (Baldwin *et al*., 2005). The coronary lymphatic vasculature of *Vegfd* mutant hearts at E16.5 showed no significant alterations (Figure 3D-K), although on the ventral side of the ventricles lymphatic vessel coverage, total lymphatic vessel length and total bifurcations showed a tendency to reduction.

**Figure 3.**
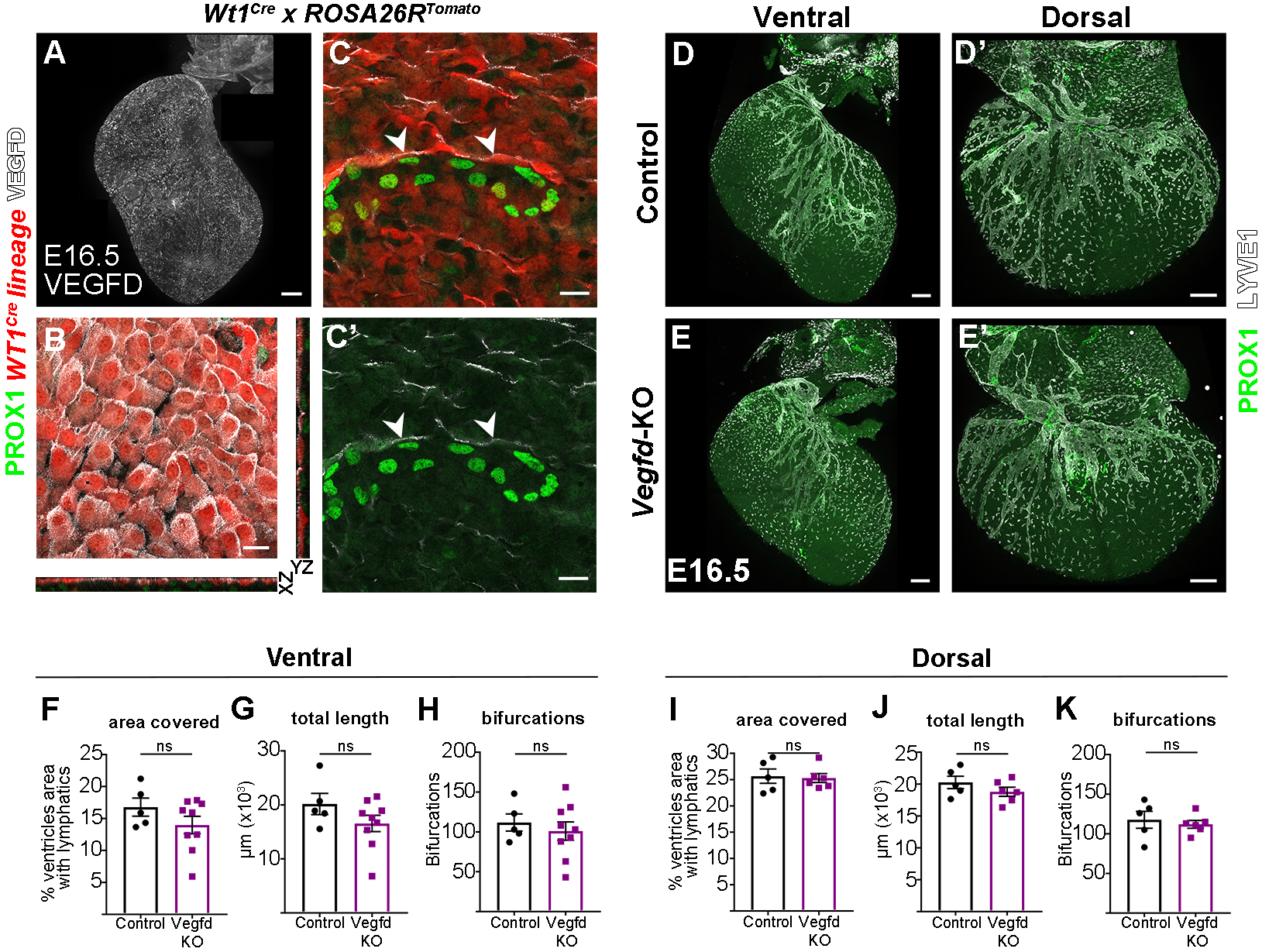
Epicardial expression of VEGFD is not essential for coronary lymphangiogenesis. **A**, confocal detection of VEGFD protein in a whole-mount E16.5 mouse heart (ventral side). **B**, Maximum projection of three epicardial confocal planes showing detection of VEGFD in the epicardium of a E16.5 *Wt1^Cre^; Rosa26R^Tomato^* mouse heart, in which the *Wt1^Cre^* lineage reveals the epicardium in red. (**C-C’**) Maximum projection of three subepicardial confocal planes showing Prox1^+^ lymphatic endothelial cells surrounded by VEGFD-expressing epicardial-derived cells (arrowheads). (**D-E’**) PROX1 and LYVE1 whole-mount immunostaining of E16.5 control and *Vegfd^-/-^ (Vegfd-KO*) hearts. (**F-K**) Quantification of the percentage of area covered by lymphatic vessels in different cardiac regions (**F, I**), total lymphatic length (**G**, **J**) and number of bifurcations (**H**, **K**) in control and *Vegfd-KO* hearts. Each point represents a quantified heart. Statistics: Mann-Whitney test with two-tailed p-values.

To determine whether epicardial VEGFD and VEGFC act redundantly in cardiac lymphangiogenesis, we studied the coronary lymphatic vasculature of compound mutants (Figure 4A-D’). We studied *Wt1^Cre^-mediated* epicardial elimination of *Vefgc* with or without elimination of *Vegfd*. Due to the location of *Vegfd* in the X chromosome, male KOs are hemizygous, while females are homozygous for the *Vegfd* deletion. Elimination of *Vegfd* in females exacerbated the reduction in coronary lymphatic vasculature observed in the epicardial deletion of *Vegfc* (Figure 4A-C’, E, F). This reduction was significant only in the ventral side of the ventricles, suggesting a stronger interaction between VEGFC and VEGFD in this part of the heart (Figure 4E, F). Unexpectedly, we found that *Vegfd* deletion in males did not worsen the reduction in coronary lymphatics observed in the epicardial deletion of *Vegfc* (Figure 4B, D, B’, D’, E, F). These results show that epicardial VEGFC and D cooperate in coronary lymphangiogenesis in females but not in males and with a stronger intensity in the ventral side of the ventricles. Given that epicardial *Vegfc* deletion affected more the dorsal coronary lymphatic vasculature, these results suggest different engagement of VEGFC and VEGFD in dorsal versus ventral coronary lymphangiogenesis.

**Figure 4.**
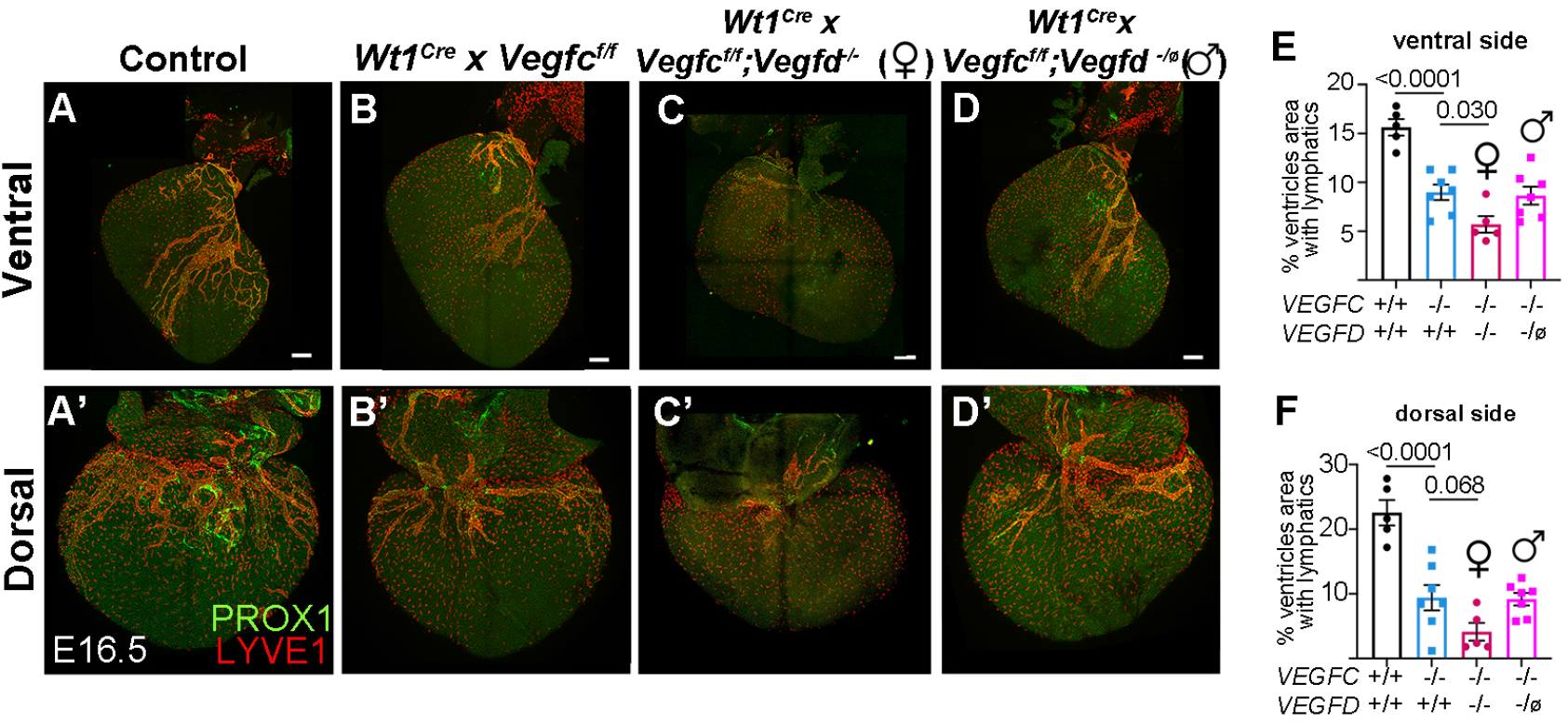
Sex-specific cooperative redundant roles of VEGFC and VEGFD in coronary lymhpangiogenesis. (**A-D’**) Representative specimens showing lymphatic vessel coverage of the dorsal and ventral ventricular surfaces upon combined conditional elimination of *Vegfc* with *Wt1^Cre^* and constitutive elimination of *Vegfd*. (**E, F**) Quantification of the ventricular lymphatic vessel coverage in combinations of *Vegfc/d* mutants. Statistics: one-way ANOVA with Sidak’s correction for multiple comparisons.

Our results thus reveal an essential role of the epicardium in coronary lymphangiogenesis through the production of VEGFC and VEGFD. Previously, elimination of epicardial VEGFC showed delayed dorsal blood coronary vasculature development (Chen *et al*., 2014), however, here we show complete agenesis of the lymphatic vasculature, which reveals regulation of lymphangiogenesis by epicardial VEGFC and VEGFD independently of the blood vasculature. While we used epicardial drivers, epicardial-derived inherit this recombination, they associate intimately with LECs, and we showed that they express VEGFD. We cannot, therefore, exclude a role of epicardial-derived cells in contributing to coronary lymphangiogenesis. Nonetheless, the epicardium appears to be the most important driver of lymphangiogenesis, given that epicardial-derived cells invade the myocardium, while lymphatic vessels always grow in intimate contact with the epicardium, without entering the myocardium. This view is in agreement with the strong expression of the lymphatic endothelial cell guidance cytokine CXCL12 in the epicardium, together with VEGFC and VEGFD (Cavallero *et al*, 2015; Chen *et al*, 2014, and this study). These findings identify the coronary lymphatic vasculature as an exception compared to other tissues and organs in which lymphangiogenesis is guided by signalling from the smooth muscle layer of pre-existing blood vessels (Cha *et al*., 2012; Vaahtomeri *et al*., 2017).

We also uncovered a sex-specific requirement for VEGFD in the heart, however, the role of VEGFD is only revealed in a defective VEGFC background, indicating that VEGFC is alone sufficient to drive coronary lymphangiogenesis in both males and females. Why males are insensitive to VEGFD is therefore difficult to characterize and may relate to sex-specific patterns of VEGFC/D expression or sensitivity to their signalling.

Finally, the different requirements of VEGFC and VEGFD in ventral versus dorsal lymphangiogenesis may relate to the recent discovery of the contribution of second heart field precursors exclusively to the ventral coronary lymphatic endothelium (Lioux *et al*., 2020). The different origins of LECs in the ventral versus the dorsal side of the ventricles may determine intrinsic differential sensitivities of the ventral and dorsal lymphatic vasculatures to VEGFC and VEGFD. In this case, the results suggest that second heart field-derived LECs could be more sensitive to the loss of VEGFD than those derived from the cardinal vein.

In summary, we report a cardiac-specific mechanism of lymphatic vasculature development that relies on essential and redundant contribution of epicardial VEGFC and VEGFD to coronary lymphangiogenesis.

## Methods

### Mouse lines

Animals were handled in accordance with CNIC Ethics Committee, Spanish laws and the EU Directive 2010/63/EU for the use of animals in research. All mouse experiments were approved by the CNIC and Universidad Autónoma de Madrid Committees for “Ética y Bienestar Animal” and the area of “Protección Animal” of the Community of Madrid with references PROEX 220/15 and PROEX 144.1/21 For this study, mice were maintained on a mixed background. Mouse experiments performed at Northwestern University were performed in accordance with protocols approved by Northwestern University Institutional Animal Care and Use Committee. The mouse alleles used here and already described were: *Wt1^Cre^* (Wessels *et al*, 2012), *Tbx18^Cre^* (Cai *et al*, 2008), *Rosa26^tdtmt^* (Madisen *et al*, 2010), *Vegfc^flox^* (Lim *et al*, 2019), *Eef1a1^VegfcGOF^* (Pichol-Thievend *et al*., 2018).

A new *Vegfd knockout* line was generated using CRISPR-Cas9 technology. Four sgRNAs that recognized *Vegfd* genomic sequences were designed using the CRISPOR web tool (Concordet & Haeussler, 2018): two sgRNAS to target intron 2 (A and B), and two to target intron 4 (C and D). sgRNA-coding sequences A) TAGGTTAAGTTCCCATATAGTGG; B)

GCGTCATGAAAAGCATGTCAGGG; C) ATGCCTGTATAATGGGTAAAGG; D) GTGCAACACATGTCTTTCTG AGG. The four sgRNAs were used simultaneously. Around 3.4 kb of *Vegfd* gene from intron 2 to intron 4 and including exon 3, intron 3 and exon 4 were deleted. Exons 3-4 code for VEGFD amino acids 106 to 218. To generate mouse mutants carrying the deletion ten 3 to 5-weeks old C57BL/6JCrl females were superovulated by injection of 5 IU of PMSG and 48 hours later with 5 IU of hCG. Females were then crossed with C57BL/6JCrl males and the next morning they were checked for positive plug. Fertilised zygotes were retrieved and incubated at 37°C with 5% CO_2_/5% O_2_ in Evolve-KSOM medium (Zenith Biotech ZEKS-050) for pronuclear microinjection of 1-2 pL containing 100μg of Cas9 protein (IDT Alt-R® S.p. HiFi Cas9 Nuclease V3, 1081060) and 0.305 μM of each sgRNA (A to D) (IDT Alt-R® CRISPR-Cas9 sgRNA). Injected embryos were then incubated in Evolve-KSOM medium at 37°C and 5% CO_2_/5% O_2_ overnight. The next morning, the embryos at two-cell stage were transferred to CD-1 pseudopregnant females. 5 microinjected animals were obtained at weaning and 3 of them had exons 3 and 4 deleted and were used to establish the mouse line. For genotyping, the following primers were used: To detect the deletion of Exon 3: Fw1: GTGCTATCCAGCTGTAGCCT; Rv1: CCCCTGAGCCTGTTTCTTTACT (Figure S10B); To detect the deletion of Exon 4: Fw2: GGGCAAAAATGCAGATGGTGG; Rv2: GATCCTCAAGGTTTTGGGTCCT (Figure S10C); To detect the total deletion of both, exon 3 and exon 4: Fw1: GTGCTATCCAGCTGTAGCCT; Rv2: GATCCTCAAGGTTTTGGGTCCT. The specific deletions obtained were characterized by Sanger DNA sequencing. We confirmed that the mutant sequences aligned with the flanking regions of *Vegfd* exons 3 and 4 and contained deletions from 276 bp upstream exon 3 to 67 bp downstream exon 4 (2 founders) or from 272 bp upstream exon 3 to 90 bp downstream exon 4 (1 founder). Females were maintained in heterozygosity and the males in hemizygosity. These animals were crossed and maintained in a C57BL/6 background.

Except for *Vegfd*-KO line, mice were genotyped by PCR as described in the original reports. Male and female mice older than 8 weeks of age were used for mating. Experimental specimens were retrieved during gestation and sex-determined by PCR for *Vegfd*-KO and *Vegfd*-KO;*Vegfc*-KO embryos.

### Embryo and Organ Retrieval

The morning in which the vaginal plug was detected was considered as embryonic day 0.5 (E0.5). Pregnant females were sacrificed by CO_2_ inhalation followed by cervical dislocation. Embryos were dissected in PBS with a sprinkle of Heparin (ROVI 1000 IU/mL). Fetuses were decapitated and placed in a 50 mM KCl (Sigma P9541) solution in PBS with heparin to stop the hearts in diastole and avoid clothing before dissection. Hearts were dissected and fixed in PFA 4% in PBS overnight at 2°C. The tip of the tail was used to genotype specimens.

### Tissue sectioning and immunostaining

Hearts were washed in PBS after fixation and cryoprotected in 15% sucrose (Sigma, 16104) PBS overnight at 4°C. The next morning, sucrose was removed and a 37°C pre-heated solution of 7.5% gelatin (Sigma, G2500), 15% sucrose in PBS was added. Hearts were incubated in this solution for at least 4 hours and then allowed to solidify at 4 °C overnight. Gelatin blocks were snap-frozen in a −70°C solution of isopentane (Sigma, 1060561000) for 1 minute. The frozen blocks were kept at −80°C until sectioned. 8 μm-thick cryo-sections were made using a Leica CM1950 automated Cryostat and stored at −20°C until used. Slides were thaw at room temperature and gelatin was removed from the slides by two 10-minute washes with PBS at 37°C and a quick wash with room-temperature (RT) PBS. For HRP-immunohistochemistry, a peroxidase quenching step was performed. By incubating at RT in darkness for 1 hour in a 1% hydrogen peroxide (Sigma, H1009) solution in 40 % methanol in PBS. Sections were then permeabilised with 0.5% Triton X-100 (Sigma T9284) in PBS for 30 minutes at RT, followed by PBS washing and treated with TNB blocking reagent (Perkin Elmer, FP1012) for 1 hour at RT. The sections were then incubated with primary-antibody dilutions prepared in the same TNB blocking solution within a humid chamber at 4°C overnight. Secondary antibodies were incubated for 1 hour at RT. Slides were washed with 0.01% Tween-20 (Sigma P9416) in PBS several times between the previous steps. Slides were mounted with Dako fluorescence mounting medium (s3023).

### Antibodies

CX-40: Alpha Diagnostics CX40-A, Rabbit 1:200; EMCN eFluor660 eBioscience^™^ 50-5851-82, Rat 1:100/1:200; LYVE1 ReliaTech 03-PA50S, Rabbit 1:200; PROX1 R&D systems AF2727, Goat 1:200; VEGFD Cusabio CSB-PA07554A0Rb, Rabbit 1:200.

### Whole-mount immunofluorescence

For hearts, after fixation and subsequent wash with PBS, all the following steps were performed at 4°C in a rotating wheel. 500 μL were added at each step, except for the washes that were performed in a 50 ml Falcon tube. Protocol: Permeabilization for 2 days in 0.5% Triton X-100 in PBS was followed by 2-hour wash with PBS. TNB blocking was performed over day and primary antibody incubation for 3 days followed by several washes overday with 0.01% Tween 20 in PBS. Secondary antibodies were incubated from overnight to 2 days, depending on the antigen and cardiac stage. Finally, hearts were thoroughly washed with 0.01% Tween 20 in PBS followed by PBS to remove detergent. To improve image acquisition while preserving fluorescence, hearts were mildly clarified with increasing glycerol concentrations starting at 20% glycerol in PBS up to 80%. For immunofluorescences of Figure 2, the blocking solution was 3% BSA, 0.1% Triton and 5% donkey serum in PBS.

### Image acquisition and quantification

Whole-mount and cardiac sections immunofluorescence were acquired with: Nikon A1R confocal microscope, Leica SP5 multiline inverted confocal microscope, Leica TCS SP8 coupled to a DMi8 inverted confocal microscope with Navigator module equipped with white light laser and Nikon W1 Spinning Disk inverted confocal microscope. Images were analysed and quantifications were made using ImageJ (http://rsb.info.nih.gov/ij). Maximum projections were acquired using TileScan and z-stack functions. The Maximum Projection of equivalent z-stacks for the different hearts was used for quantification. We used the Freehand selection tool of ImageJ to select the whole ventral or dorsal ventricular surfaces and the area covered by Lymphatic vessels. For LYVE1 whole-mount quantifications of Vegfd-KOs, Vegfc-GOF and VEGFC-KO;VEGFD-KO hearts, a macro was used to quantify lymphatic coverage. This included several steps: duplicate channel of interest, median filter, binary conversion to Mask, analyse particles tool to remove smaller particles (macrophages) and finally, after creating a raw selection, manual curation using the Freehand selection tool. To quantify the total lymphatic length the Freehand line tool was used to draw lines running medial along each lymphatic branch. Total length was obtained by adding the length of every branch. Bifurcations were manually counted, considering each intersection point of two or more lymphatic branches.

### Statistics

The details of each test used are specified in Figure legends. As a general rule, Mann-Whitney two-tailed test was performed to compare two groups of quantitative data. One-way ANOVA was performed to compare more than two groups of quantitative data with one independent variable, after a normality test (Kolmogorov-Smirnov) was found positive for all samples. Data is indicated as mean ± SEM of the individual plotted values. All comparisons and graphs were made using GraphPad Prism 9 statistical analysis software. In all cases p-values were two-tailed and adjusted by Sidak’s correction for multiple measurements. Values of p ≥ 0.05 were considered non-significant.

## Data Availability

All source data will be made available from the Mendeley public repository.

## Acknowledgements

We thank Cristina Villa and the Torres group for helpful comments and discussions, Xiaolei Liu and Michael Oxendine for advice during experimentation, Wanshu Ma for the *Eef1a1^VegfcGOF^* mouse strain, Wanshu Ma and Mark Kahn for the *Vegfc^flox/flox^* mouse, and the CNIC Genomics, Microscopy, Cellomics and Transgenesis Units personnel for their support to this work. This work was supported by the European Commission H2020 Program grant SC1-BHC-07-2019. Ref. 874764 “REANIMA” to M.T.; the Spanish Ministerio de Ciencia e Innovación grant PGC2018-096486-B-I00 to M.T.; Grant TNE-17CVD04 from the Leducq Foundation to M.T.; Comunidad de Madrid grant P2022/BMD-7245 to M.T.; RO1HL151388 and RO1HL162800 to GO, FPU grant from the Spanish Ministry of Education, Culture and Sports (Grant FPU15/02955) and EMBO Short-Term Fellowship number 8357 to E.d.C.; for experiments in the Unidad de Microscopía e Imagen Dinámica, CNIC, ICTS-ReDib, MCIN/AEI /10.13039/501100011033 and FEDER “Una manera de hacer Europa” (#ICTS-2018-04-CNIC-16). The CNIC is supported by the Ministerio de Ciencia e Innovación and the Pro CNIC Foundation, and is a Severo Ochoa Center of Excellence (Grant number CEX2020-001041-S).

## Author Contributions

E.d.C., Conceptualization, Methodology, Formal Analysis, Investigation, Visualization, Writing-Original Draft, Writing-Review and Editing, Funding Acquisition. S.T. and V. C., Methodology; G.O., Supervision, Resources, Writing-Review and Editing, Funding Acquisition. M.T. Conceptualization, Writing-Review and Editing, Supervision, Funding Acquisition.

